# Pharmaceutical efficacy of human epiphyseal chondrocytes with differential replication numbers for cellular therapy products

**DOI:** 10.1101/085688

**Authors:** Michiyo Nasu, Shinichiro Takayama, Akihiro Umezawa

## Abstract

The cell-based therapy for cartilage or bone requires a large number of cells serial passages of chondrocytes are, therefore, needed. However, fates of expanded chondrocytes from extra fingers remains unclarified. The chondrocytes from human epiphyses morphologically changed from small polygonal cells, to bipolar elongated spindle cells and to large polygonal cells with degeneration at early passages. Gene of type II collagen was expressed in the cells only at a primary culture (Passage 0) and Passage 1 (P1) cells. The nodules by implantation of P0 to P8 cells were composed of cartilage and perichondrium. The cartilage consisted of chondrocytes with round nuclei and type II collagen-positive matrix, and the perichondrium consisted of spindle cells with type I collage-positive matrix. The cartilage and perichondrium developed to bone with marrow cavity through enchondral ossification. Chondrogenesis and osteogenesis by epiphyseal chondrocytes depended on replication number in culture. It is noteworthy to take population doubling level in correlation with pharmaceutical efficacy into consideration when we use chondrocytes for cell-based therapies.

## Introduction

Epiphyseal cartilage of long bone is destined to develop to bone with marrow cavity and articular cartilage. The epiphyseal cartilage continues to proliferate and undergoes enchondral ossification until apparent growth ceases at adolescence, and eventually is replaced with bone except for articular surface which remains cartilaginous throughout life [1]. Chondrocytes in culture have an ability to generate cartilage in vivo, and can be a tool to simulate developmental process. Our previous study demonstrated that human infant epiphyseal chondrocytes can also simulate developmental process of epiphyseal cartilage as well as other cell types [2, 3]. Chondrocytes in culture regenerate the cartilaginous tissue by a short-term implantation in vivo and exhibited enchondral ossification after a longer period. The cell-based therapeutic strategies in cartilage and bone require serial cell passage of expanded chondrocytes for acquisition of a large number of cells. In this study, we extensively characterized chondrocytes derived from human epiphyseal cartilage in correlation with regenerative medicine.

## Materials and Methods

### Human epiphyseal chondrocytes (HECs)

HECs were obtained from epiphyseal cartilage of the phalanx and metacarpal bones of amputated supernumerary digitus with polydactyly of finger or toe of 5 patients (patient’s age 7-14 months). Signed informed consents were obtained from the parents of the donors. Isolation of HECs was performed as described previously [2]. Briefly, after confirmation of no development of secondary ossification center in the epiphyseal ends of bone, the epiphyseal cartilage was dissected out from bones and cut into piece of 1 mm^3^. The diced fragments were placed in 0.2% trypsin/0.02% EDTA solution (Immuno-Biological Laboratories Co, Ltd. Gunma, Japan) diluted with PBS at 37°C for 30 min and incubated in Dulbecco’s modified Eagle’s medium (DMEM, Sigma-Aldrich Japan K.K. Tokyo) with 0.1% collagenase (Wako Pure Chemical Industries, Ltd. Osaka,Japan) at 37°C for 7 to 10 h. After centrifuge of the isolated cells, the cell-pellets were resuspended in DMEM. Isolated chondrocytes were grown in MF basal medium with 1% fetal calf serum (Toyobo Life science, Fukui, Japan) at a density of 0.5-1 x 10^5^ cells / 100-mm dish and 10-mm chamber slide by a monolayer culture in 37°C and 5% CO_2_ incubator.

### Cytology

Culture cells on the chamber slides were stained by Papanicolaou’s method after fixation in 100% ethanol and by senescence-associated ß-galactosidase [4].

### Reverse transcription polymerase chain reaction (RT-PCR)

Total RNA was isolated from HECs, using the RNeasy kit (Qiagen, Valencia, Ca, USA).Total RNA (2 µg) was reverse-transcribed to cDNA at 50°C for 50 min in a volume of 20 µl containing the following reagents: 0.5 mM dNTP mix, 0.5 µM oligo-dT_12-18_ primer, first strand buffer, RNase inhibitor and 200 unit Superscript III (RNase H-free reverse transcriptase). All the reagents were purchased from Invitrogen (Carlsbad, CA,USA). After terminating the reaction at 70°C for 15 min, 1 U RNase H (Invitrogen) was added to the reaction mixture, followed by incubation at 37°C for 10 min to remove RNA. PCR was performed in a total volume of 50 µl, using the primers (Table 1) and the Ex Taq Hot Start version (Takara Bio Inc. Shiga, Japan). The PCR products were electrophoresed, and visualized on ethidium bromide-stained 1.5% agarose gels.

**Table 1.**
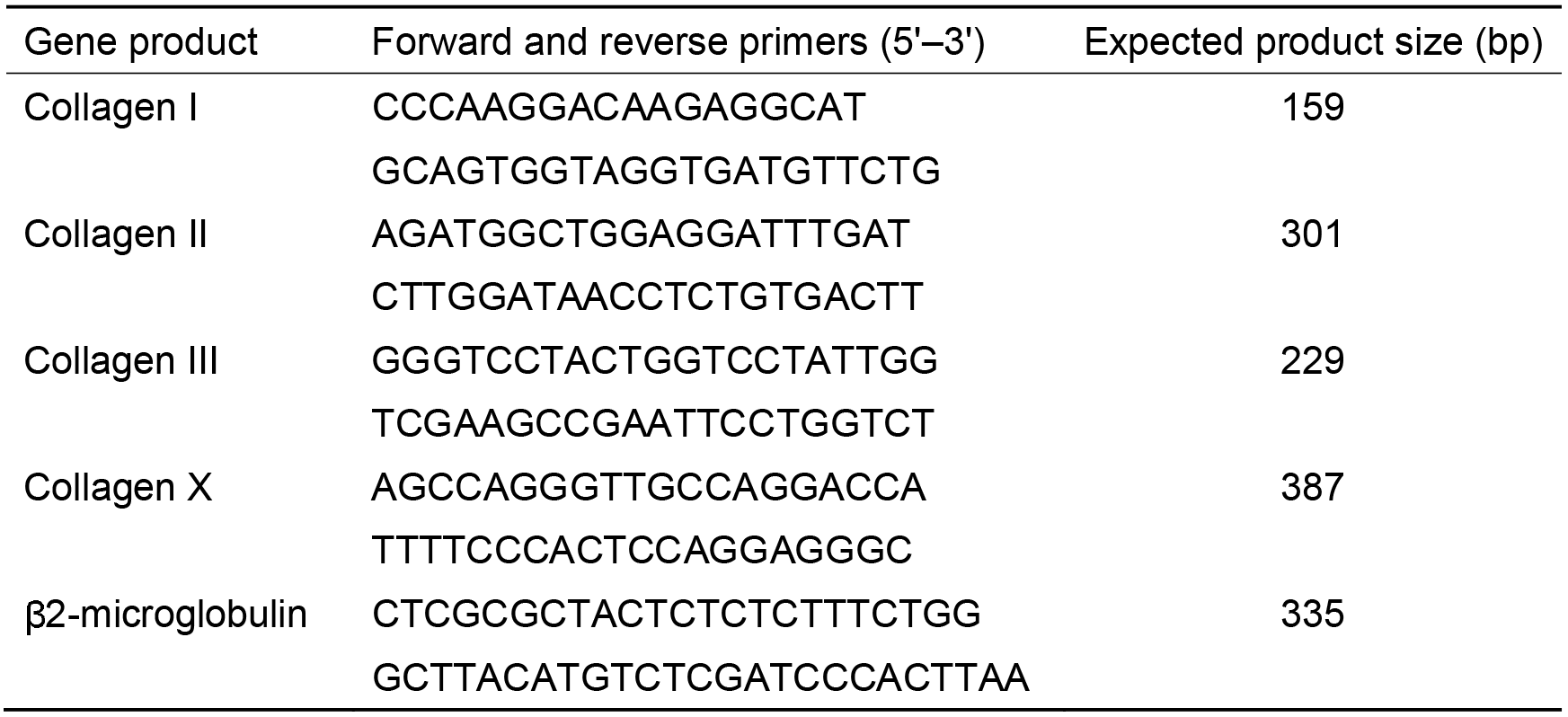
Primer pairs and experimental conditions for RT-PCR

### Cell implantation

To determine the ability of HECs in a monolayer culture, P0 to P8 cells of 1.5-3 x 10^7^ in number were subcutaneously inoculated into the back skin of 6 to 8 week old,NOD/Shi-*scid*, IL-2Rγ^null^(NOG) mice (Central Institute for Experimental Animals,Kanagawa, Japan). The subcutaneous nodules of the mice were removed, and photographed. Their weight and size were then measured.

### Microscopic analysis with immunohistochemistry

Subcutaneous nodules in mouse backs were fixed in 4% paraformaldehyde and embedded in paraffin, after decalcification with 5% EDTA (pH 6.8) solution if necessary. Cutting sections were deparaffinized, dehydrated, and stained with hematoxyline and eosion (HE), and with toluidine blue (pH 7.0) and alcian blue (pH 2.5) for cartilaginous matrix.

For immunohistochemical analysis, the following primary antibodies were used: mouse monoclonal anti-human collagen type II (clone: II-4c11) and type I (clone: I-8H5) antibododies (Daiichi Fine Chemical Co. Ltd, Toyama, Japan). Additionally, anti-human antibodies (Dako, Glostand, Denmark) of vimentin (clone: V9), CD31 (clone: JC70A) and CD34 (clone: QBEnd 10) were used. These antibodies do not react with mouse antigens according to applications for use, and furthermore these things have been reconfirmed in our previous experiment [2]. Cut paraffin sections were deparaffinized, dehydrated and treated with 2% proteinase K (Dako) in Tris-HCl buffer pH 7.5 solution for 5 min at room temperature, or heated in ChemMate Target Retried Solution (Dako) for 5-20 min in a high-pressure steam sterilizer for epitope unmasking. After blocking of endogenous peroxidase by 1% hydrogen peroxide/methanol for 15 min, the sections were incubated at room temperature for 60 min in primary antibodies diluted with antibody diluent (Dako). The sections were washed with 0.01 M Tris buffered saline (TBS) solution (pH 7.4) and incubated with goat anti-mouse and anti-rabbit immunoglobulin labeled with dextran molecules-horseradish peroxidase (EnVision, Dako) for 30 min at room temperature. After washing with TBS, the sections were incubated in 3,3’-diaminobenzidin in substrate-chromogen solution (Dako) for 5-10 min.Negative controls were performed by omitting the primary antibody. The sections were counterstained with hematoxylin.

## Results

### HEC in culture

HECs were fibroblast-like in morphology at a primary culture. HECs slowly proliferated until 4th day after start of the cultivation, and then rapidly proliferated at 5-6th day (Figure 1A). The cells reached to senescence before PDL 60. PDLs of the senesced cells were 52.4 in an average at Passage 8. We then examined HECs for expression of cartilage-associated genes at each passage by RT-PCR analysis (Figure 1B). The type-II collagen gene was expressed at P0 and P1, but was not detected at P2 to P8. The genes for collagen type I, III and X remained to be expressed from P0 to P8.

**Figure 1.**
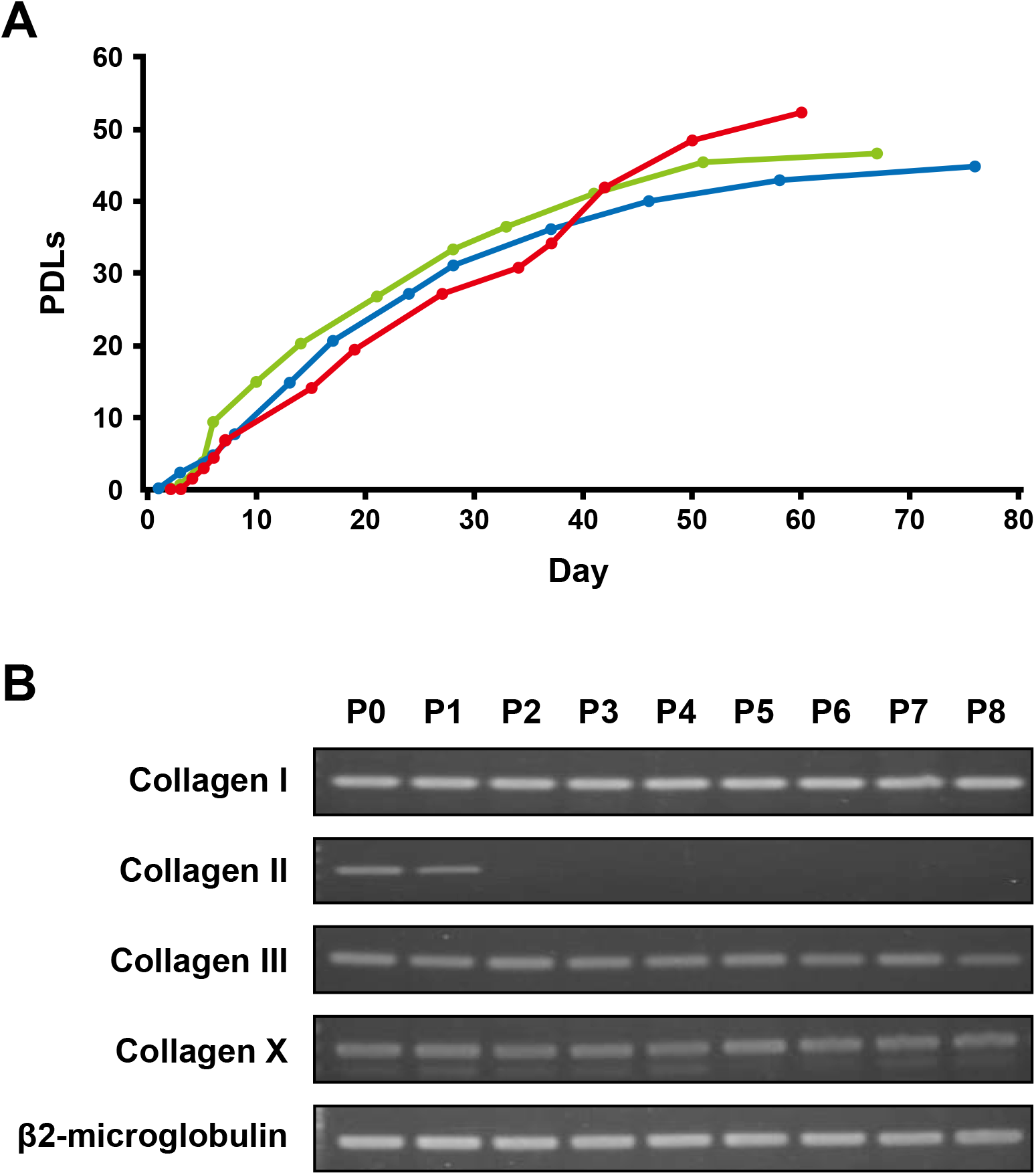
Characteristics of the human epiphyseal chondrocytes (HECs) A. Proliferative capacity of HECS. The number of cells was counted at each passage.Total number of population doubling levels (PDLs) was calculated, using the formula of log_10_ (total number of cells/starting number of cells)/log_10_2. B. RT-PCR analysis for cartilage-associated genes. P: Passage number.

HECs showed extensive morphological alternations during passage expansions (Figure 2): P0, small polygonal cells with many mitoses; P1 and P2, bipolar elongated spindle cells; P3-P5, bipolar short spindle cells of with appearance of large cells; P6-P8, large,fusiform or polygonal cells with long cytoplasmic processes, 1 indistinct nuclear 2 chromatin structures and occasional vacuolations. P8 cells became enlarged and deeply 3 stained with senescence-associated ß-galactosidase.

**Figure 2.**
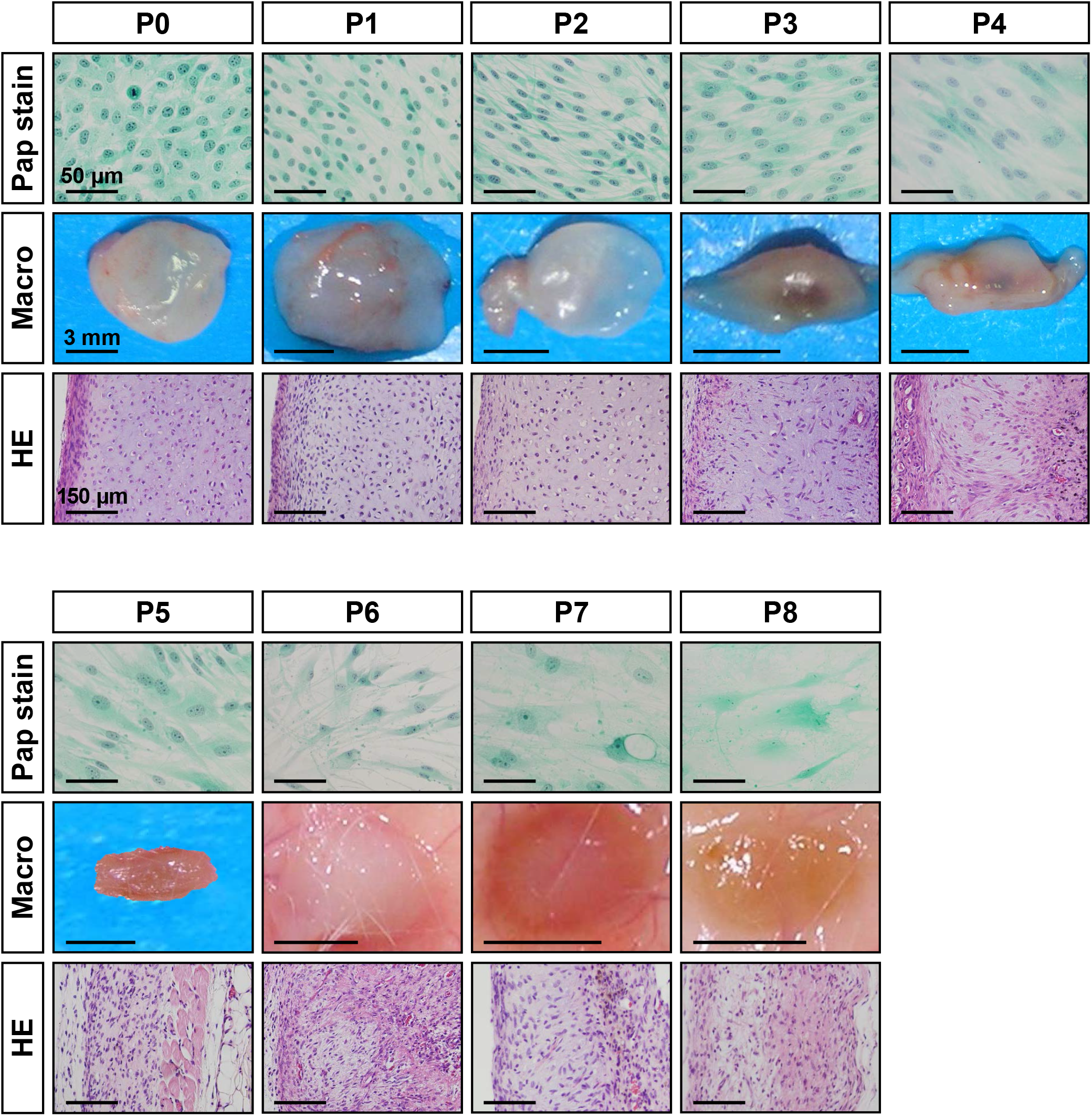
Morphological changes from P0 (PDL 8) to P8 (PDL 45) cells of human epiphyseal chondrocytes during expansion. **Top panels**: Cytology of the culture cells with Papanicolaou’s stain (Pap stain). Cell morphology changed from small polygonal cells with mitosis (P0) to short or long (P1),to long spindle cells (P2-P4) with occasional large spindle or polygonal cells (P4-P5)and to large, polygonal cells with long cytoplasmic processes and vacuolation, and nuclei with indistinct chromatin structures (P6-P8). **Middle panels**: Macroscopic views of the tissues by implantation of P0 to P8 cells for two weeks. The nodules of P0 to P2 cells are the same in size, in weight and in degree of transparency with whitish color on surface. The nodule of P3 and P4 cells, however, decreased in size along with the increase of passages, and were accompanied central vascular invasion. The nodules of P5 to P8 cells became flattened and to be white to brown in color. Bars: 3 mm. **Bottom panels**: Microscopic views with hematoxylin and eosin stain. P0 and P1 cells generated two distinct portions, i.e. cartilage and perichondrium at the implanted sites.As the increase in passage number, the boundary of two portions became to be ill-defined. P4 to P8 cells exhibited spindle-shaped dedifferentiated chondrocytes, accompanied with reactions of inflammatory cells, macrophages and foreign-body giant cells.

### Implantation of HECs for two weeks

We subcutaneously implanted HECs at each passage to investigate ability of chondrogenesis and osteogenesis in vivo. P0 and P1 cells generated the nodules that hardly changed in size, weight and in degree of transparency with whitish color on surface (Figure 2). In contrast, P2 cells foamed the nodules that decreased in weight, size and degree of transparency. The nodule by P0 cells was clearly divided into two different parts, cartilage composing of chondrocytes with lacunae and abundant extracellular matrix in the center, and perichondrium with spindle cells-layers at the periphery (Figure 3), as demonstrated in the previous study [2]. Extracellular matrix in the cartilage generated by P0 and P1 cells demonstrated metachromasia with Toluidine Blue stain. Vimentin was detected in both chondrocytes and perichondral spindle cells. Type II collagen-positive matrix was diffusely distributed in the cartilage by P0 and P1 cells, and type I collagen-positive matrix was localized in the perichondrium.

**Figure 3.**
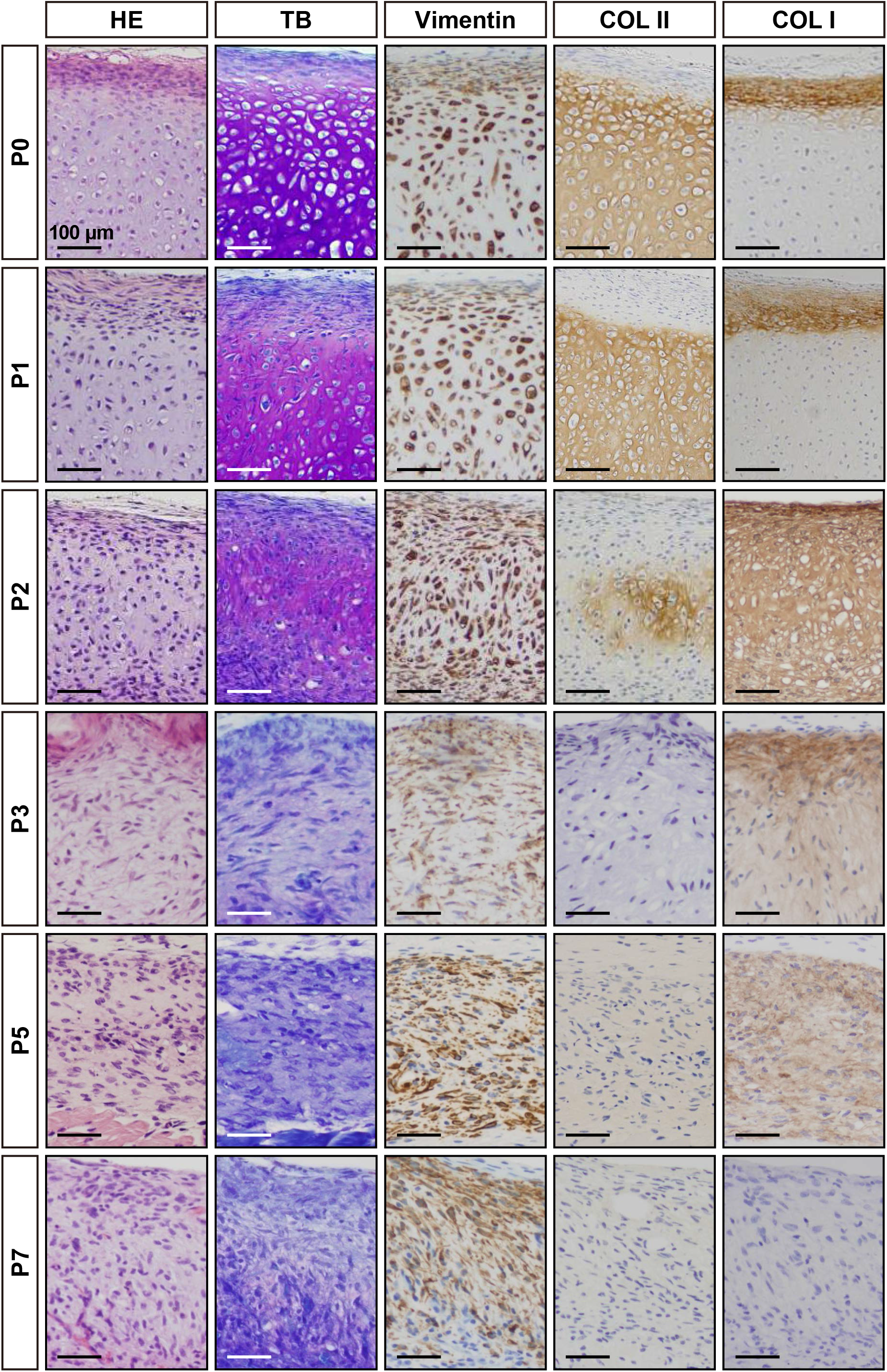
Histological and immunohistochemical analysis of the implanted sites. Hematoxylin and eosin (HE) stain, Toluidine blue (TB) stain, and immunohistochemical analysis by using antibodies against vimentin, type II collagen (COL II), and type II collagen (COL II) are shown from left to right. With TB stain, the extracellular matrix of cartilage exhibited metachromasie diffusely in the nodules by P0 and P1 cells, and focally in the nodules by P2 cells. The matrix of the nodules by P0 and P1 cells was positive for COL II at the center and with type I collagen (COL I) at the periphery. The nodule by P2 cells was focally COL II-positive and diffusely COL I-positive matrix. The matrix in the nodules by P3 and P5 cells was not stained with COL II and showed weakly positive reaction with COL I. The nodules by P7 cells were negative for COL II and COL I. The microscopic views (the same as Figure 2C) with HE stain (HE) are also shown for reference.

Unlike P0, P1 and P2 cells, P3 to P8 cells generated small and flattened nodules without clear vascular invasion on opaque, whitish to brownish surface color, and failed to generate cartilage. Microscopically, the nodules showed fibrous tissues with lack of chondrocytes with lacunae, and the increase of spindle and dedifferentiated chondrocytes. The fibrous tissue included inflammatory cells, reactions of foreign-body giant cells and vascular invasions.

### Long-term observation of cartilage and bone by P0 to P2 HEC implantation

We then followed up the nodule of bone and cartilage following implantation of HEC at P0 (PDL 7) until 20 and 40 weeks (Figure 4). Macroscopically, the nodules showed smooth surface with white-yellow and reddish portion in color at 20 weeks and 40 weeks. Enchondral ossification progressed in the cartilage at 20 weeks and new bone and marrow cavity were formed (Figure 4A-G). Bone marrow cavity was surrounded by osteoid and new bone with osteoblasts, osteoclasts, immature adipocytes, endothelial cells and fibroblastoid cells. The nodules with bone and cartilage increased in size and weight, became reddish on surface and developed prominent vascular networks at 40 weeks (Figure 4H-Q). The nodules resembling articular cartilage and cortical bone with periosteum demonstrated the thin trabecular and spicular networks of the lamellar bone with diffuse accumulation of unilocular mature adipocytes and vascular development.

**Figure 4.**
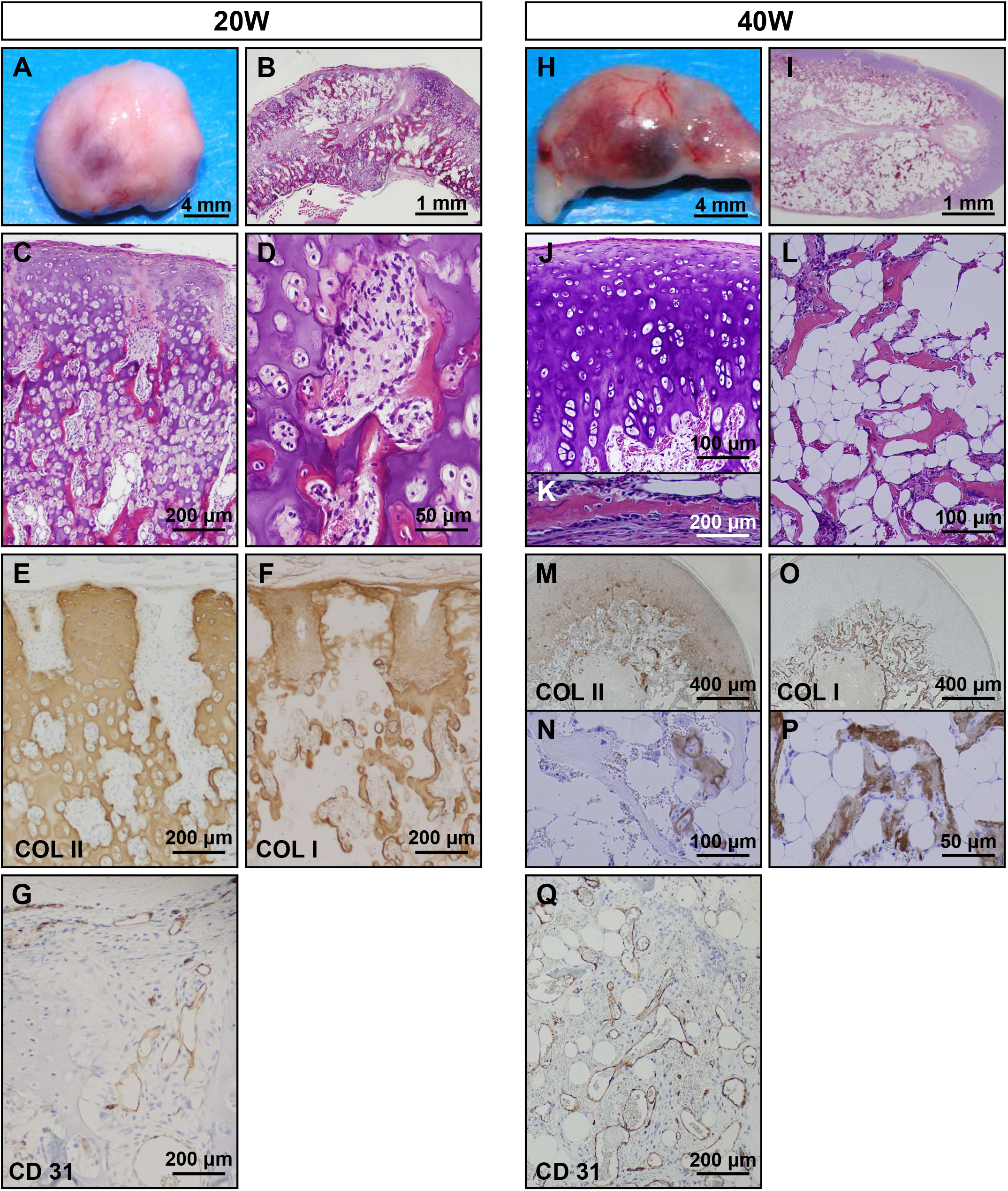
Long-term observation at the P0 cell (PDL 7)-implanted sites. P0 cell-implanted sites were analyzed at 20 weeks **(A-G)** and 40 weeks **(H-Q)**.Macroscopic examination of the P0 cell-nodules (606 mg in weight) at 20 weeks showed smooth and whitish surface with focal reddish portion and vascular networks **(A)**. Microscopic examination showed progress of enchondral ossification **(B)** with new bone formation and poor marrow cavity **(C)**. Marrow cavity consisted of osteoblasts,osteoclasts, undifferentiated mesenchyme cells and a few of immature adipocytes **(D)**. Cartilage and core of the enchondral ossification were positive for COL II **(E)**.Perichondrium, bone and osteoid was positive for COL I **(F)**. Endothelium in the early marrow cavity was positive for CD31 **(G)**. Macroscopic examination of the P0 cell-nodules (686 mg in weight) at 40 weeks showed increased reddish portion with prominent vascular networks on the surface **(H)**. Bone marrow cavity was surrounded by the cartilage and bone **(I)**,which morphologically resembled to articular cartilage **(J)**and cortical bone **(K)**, respectively. Trabecular bone was extended with the fatty marrow **(L)**. Peripheral cartilage **(M)** and core of the enchondral ossification **(N)** were positive for COL II. Perichondrium, periosteum **(O)** and trabecular bones **(P)** were positive for COL I. The vessels in the bone marrow cavity were positive for CD31 **(Q)**.

At 20 weeks, type II collagen was detected in the extracellular matrix of cartilage, but gradually reduced (Figure 4E). In contrast, type I collagen became positive in matrix of osteoid, new bone and periosteum (Figure 4F). The CD 31-positive endothelial cells were present along with the vessels at the periphery of cartilage (Figure 4G). At 40 weeks, type II collagen remained positive in peripheral cartilage and occasionally in core of enchondral ossification (Figure 4M, N). Type I collagen-positive matrix was clearly distributed in trabecular bone (Figure 4O, P). The vasculogenesis with CD 31-positive endothelial cells became prominent (Figure 4Q).

We then investigated if P1 cells are able to generate cartilage and bone like P0 cells (Figure 5). To this end, we performed 10 implantations of P1 cells (PLD 10). P1 cells under PDL 12 exhibited cartilage formation with enchondral ossification after implantation. The nodules at 25 weeks were composed of the perichondrium and cartilage with accumulation of extracellular matrix (Figure 5A, B). The cartilage was consisted of small chondrocytes with round or occasional spindle nuclei and lacunae, and covered by perichondrium with thin layer of elongated fibroblast-like spindle cells (Figure 5C). At 41 weeks (Figure 5G-L), enchondral ossification was focally demonstrated in the nodules with invasion and proliferation of peripheral undifferentiated mesenchymal cells (Figure 5H, I). Marrow-like structure with vasculogenesis, adipogenesis and osteoclastogenesis were confirmed, however, bone marrow cavity with differentiation into trabecular bone or mature adipocytes was not detected. Type I collagen-positive matrix diffusely distributed at the perichondrium at 25 weeks (Figure 5E) and at the osteoid and new bone in the enchondral ossification at 41 weeks (Figure 5K). Vasculogenesis with CD 34-positive endothelia (Figure 5F) started to be seen during enchondral ossification at 41 weeks (Figure 5L).

**Figure 5.**
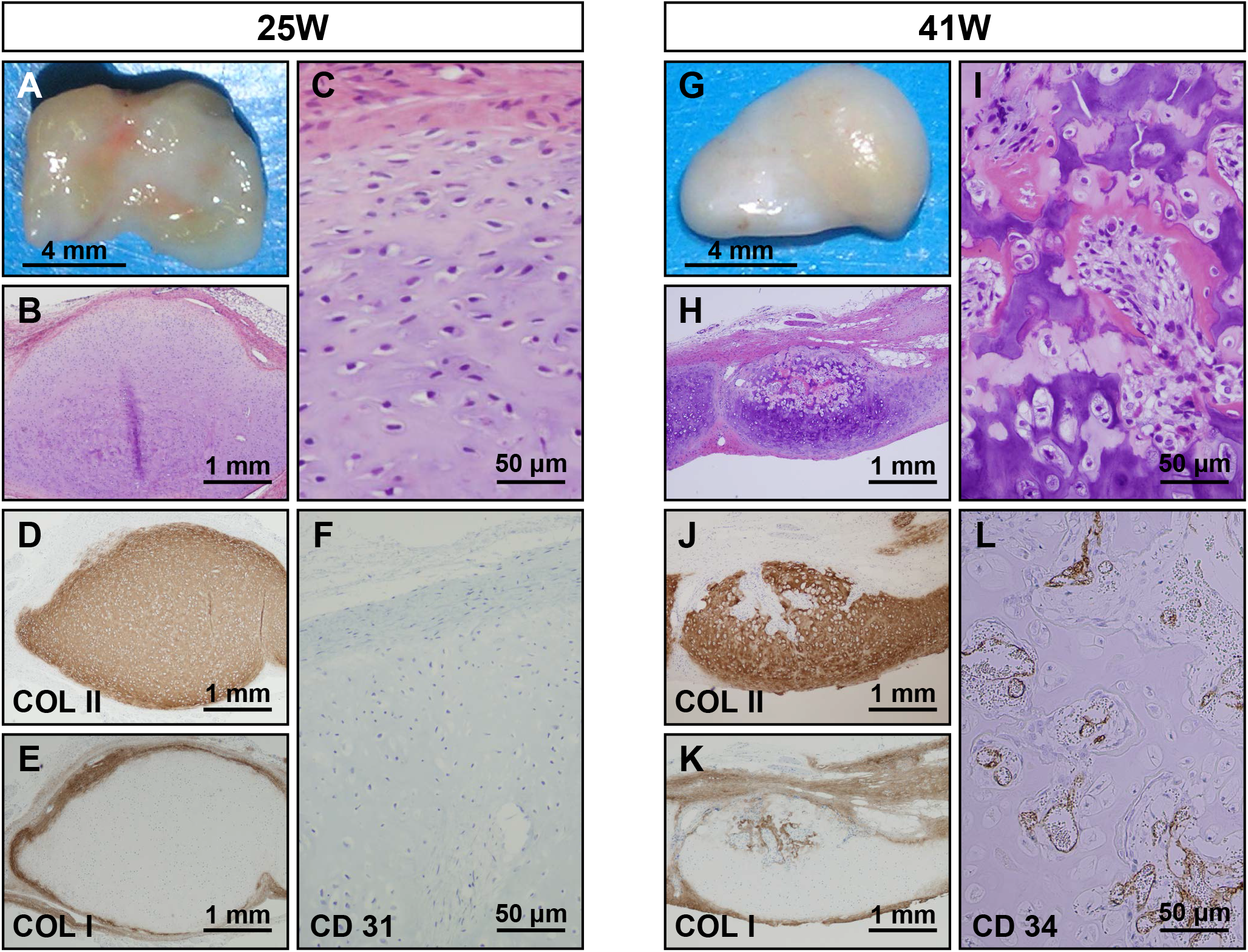
Long-term observation at the P1 cell (PDL 10)-implanted sites. P1 cell-implanted sites were analyzed at 25 weeks **(A-F)** and 41 weeks **(G-L)**. Macroscopic examination of the P1 (PDL 10) cell-nodules (86 mg in weight) at 25 weeks showed smooth and semi-translucent surface with irregular light reddish portion **(A)**. Microscopic examination showed cartilage with thin perichondrium **(B)**. The cartilage was composed of small chondrocytes with round or spindle nuclei and lacunae in the extracellular matrix **(C)**. Matrix of the cartilage **(D)** and perichondrium **(E)** were positive for COL II and COL I, respectively. CD34-positive vessels were not detected **(F)**. Macroscopic examination of the P1 cell-nodules (69 mg in weight) at 41 weeks showed smooth and whitish surface with focal yellowish portion **(G)**. Microscopic examination revealed focal enchondral ossification in the cartilage **(H)** and bone formation with early marrow cavity **(I)**. Cartilage with enchondral ossification was positive for COL II **(J)**, and perichondrium, osteoid and new bone **(K)** were positive for COL I. CD34-positive vessels were expanded in the early bone marrow cavity **(L)**.

P1 cells (PDL 14) generated cartilage that were macroscopically flattened or club-shaped, translucent and whitish surface, exhibited type II collagen-positive matrix in the cartilage at 25 and 42 weeks after the implantation, but failed to develop bone marrow even after follow-up observation (Figure 6). The cartilage at 25 weeks level was composed of small chondrocytes with round or spindle nuclei and lacunae, and hypertrophic cells with large lacunae. The perichondrium showed the structure of thick or thin layers with elongated fibroblast-like spindle cells (Figure 6B). The nodules at 42 weeks showed morphologies similar to those at 25 weeks, except decrease of hypertrophic cells with large lacunae and slight increase of chondrocyte with spindle nuclei and narrow lacunae in number, and increase in eosinophil of the perichondrial matrix and in thickness of the perichondrium. Type-II collagen-positive matrix was detected in the cartilage at 25 weeks (Figure 6D) and 42 weeks (Figure 6I). Type-I collagen-positive matrix was distributed in the thick perichondrium at 25 weeks (Figure 6E), and 42 weeks (Figure 6J).

**Figure 6.**
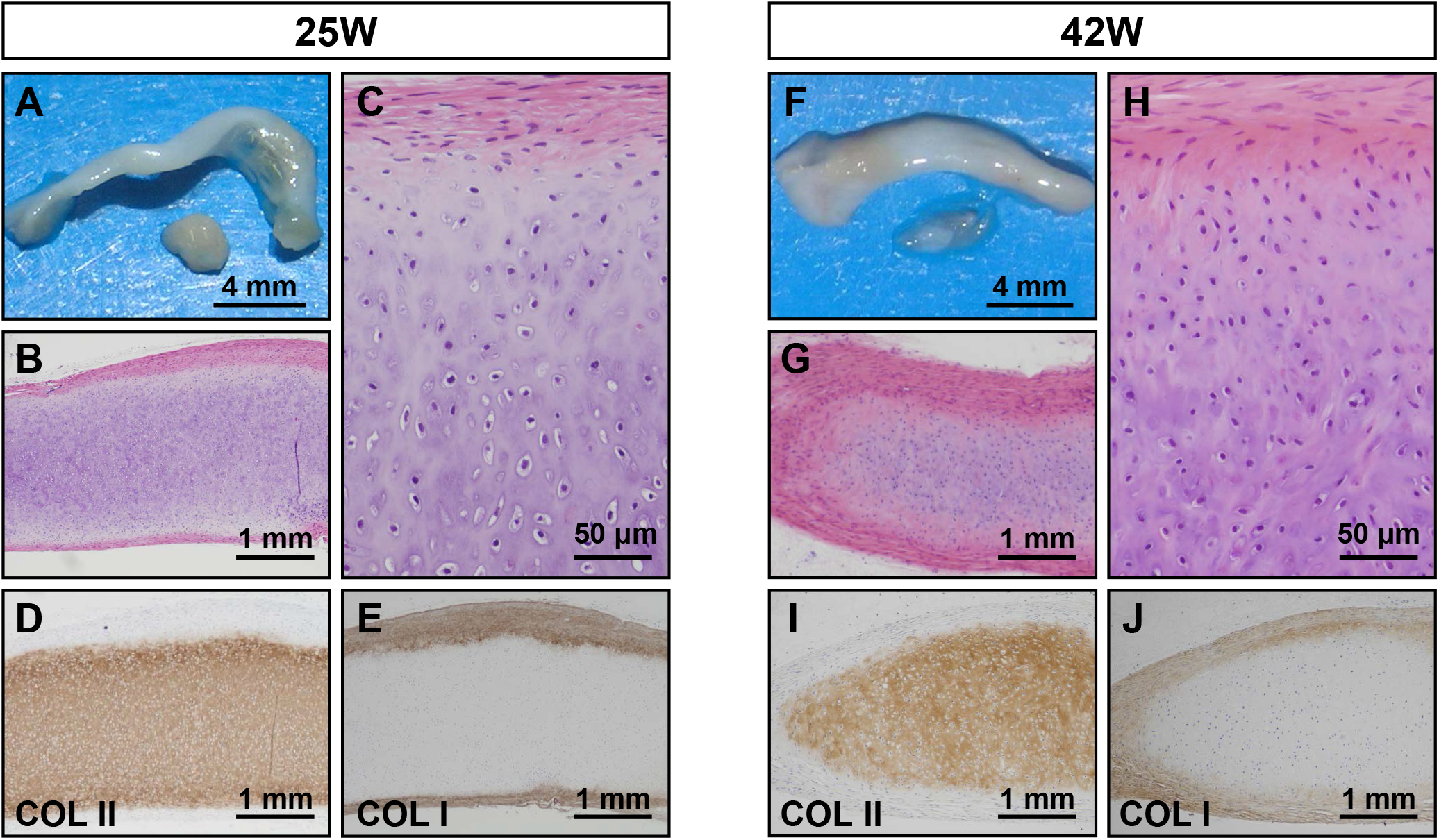
Long-term observation at the P1 cell (PDL 14)-implanted sites. P1 cell-implanted sites were analyzed at 25 weeks **(A-E)** and 41 weeks **(F-J)**. Macroscopic examination of the P1 (PDL 14) cell-nodules (73 mg in weight) at 25 weeks showed club-shaped, semi-translucent and whitish surface **(A)**. Microscopic examination showed cartilage and perichondrium **(B)**. The cartilage was composed of small chondrocytes with lacunae and of hypertrophic cells with large lacunae **(C)**. The cartilage and perichondrium were positive for COL II **(D)** and COL I **(E)**, respectively.Macroscopic examination of the P1 cell-nodules (70 mg in weight) at 42 weeks showed the same appearance as that of 25 weeks **(F)**. Microscopic examination revealed decrease of hypertrophic chondrocytes in the cartilage and increase of matrix in the perichondrium **(G, H)**. Again, the cartilage was positive for COL II **(I)** and the perichondrium was positive for COL I **(J)**.

In contrast, P2 cells also generated cartilage that was maintained for up to 17 weeks (Figure 7). Macroscopically, the cartilage showed translucent and smooth, surface that were white, partly light brown or yellow in color (Figure 7A). The cartilages at 9 weeks consisted of hypertrophic chondrocytes with round or spindle nuclei and large lacunae (Figure 7B, C). The cartilages at 17 weeks showed increase of small chondrocyte with spindle nuclei and eosinophilic extracellular matrix at the perichondrium (Figure 7G, H). Type II collagen-positive matrix remained in the cartilage at 9 weeks (Figure 7D) and at 17 weeks (Figure 7I). Type-I collagen-positive matrix was distributed in the perichondrium at 9 weeks (Figure 7E), and extended into the cartilage at 17 weeks, in addition to the perichondrium (Figure 7J).

**Figure 7.**
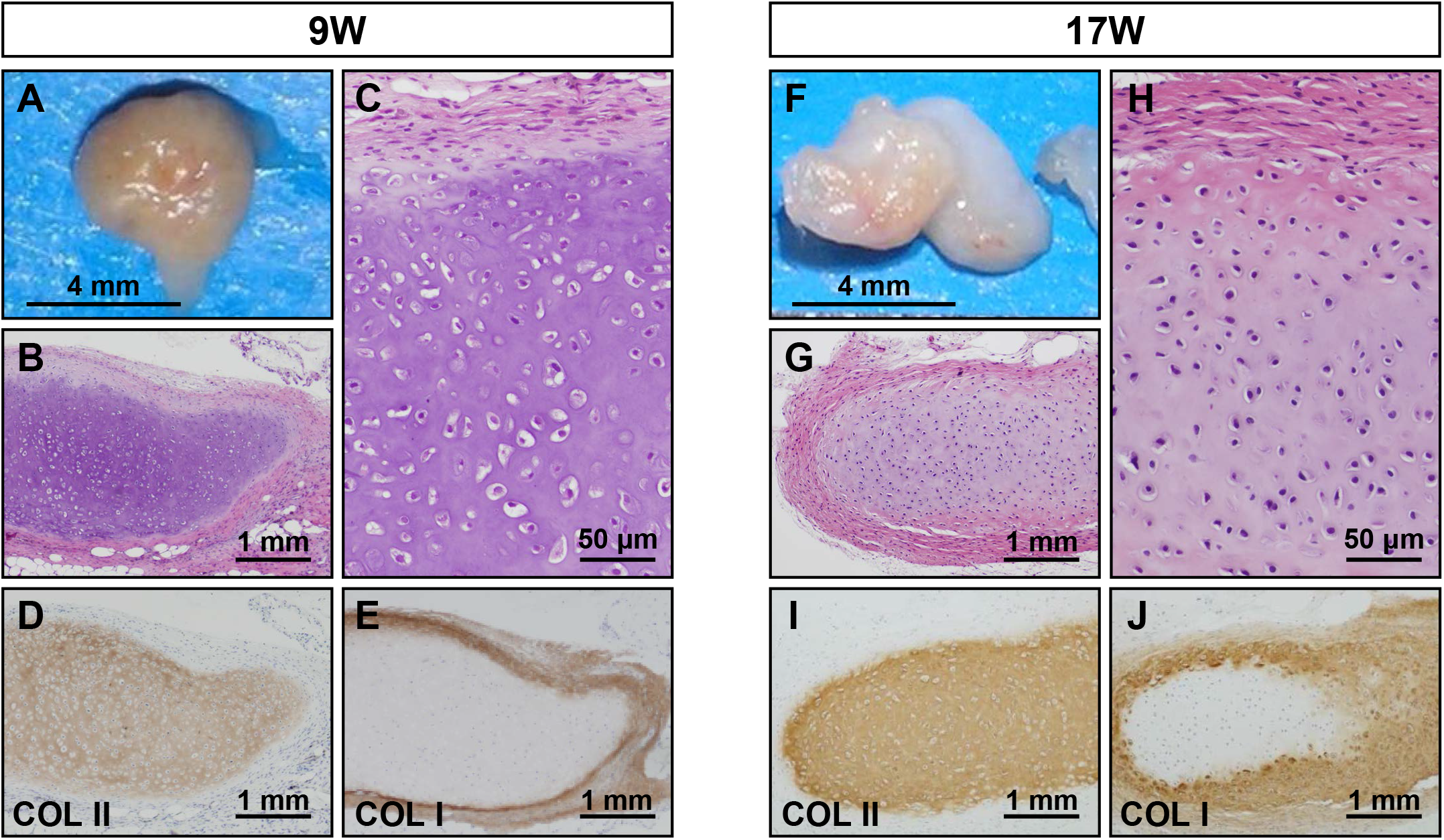
Long-term observation at the P2 cell (PDL 18)-implanted sites. P2 cell-implanted sites were analyzed at 9 weeks **(A-E)** and 17 weeks **(F-J)**. Macroscopic examination of the P2 (PDL 18) cell-nodules (9 mg in weight) at 9 weeks showed light-brown surface **(A)**. Microscopic examination showed the cartilage with small chondrocytes with round or spindle nuclei and hypertrophic cells with lacunae **(B)**and thick perichondrium **(C)**. Immunohistochemically, the cartilage was positive for COL II **(D)** and the perichondrium was positive for COL I **(E)**. Macroscopic examination of the P2 (PDL 18) cell-implanted sites (32 mg in weight) at 17 weeks showed whitish surface with partly yellowish area **(F)**. Microscopic examination showed the cartilage and irregularly thickened perichondrium **(G)** and the perichondrium with decrease of hypertrophic chondrocytes and increase of eosinophilic matrix **(H)**, compared with nodules at 9 weeks. The cartilage was positive for COL II **(I)**, and partially positive for COL I **(J)**. The perichondrium was positive for COL I.

### Lack of chondrogenic activity in P3 cells

P3 cells (PDL 23) generated fibrous tissue that was macroscopically translucent and flattened, and tightly adhered to the subcutaneous tissue (Figure 8A). The tissue was ill-defined in the boundaries between the perichondrium and cartilage (Figure 8B, C). Type II collagen-positive matrix was irregularly distributed in the cartilage (Figure 8D). Type I collagen-positive matrix was irregularly found in the cartilage and not overlapped with type II collagen-positive matrix (Figure 8E).

**Figure 8.**
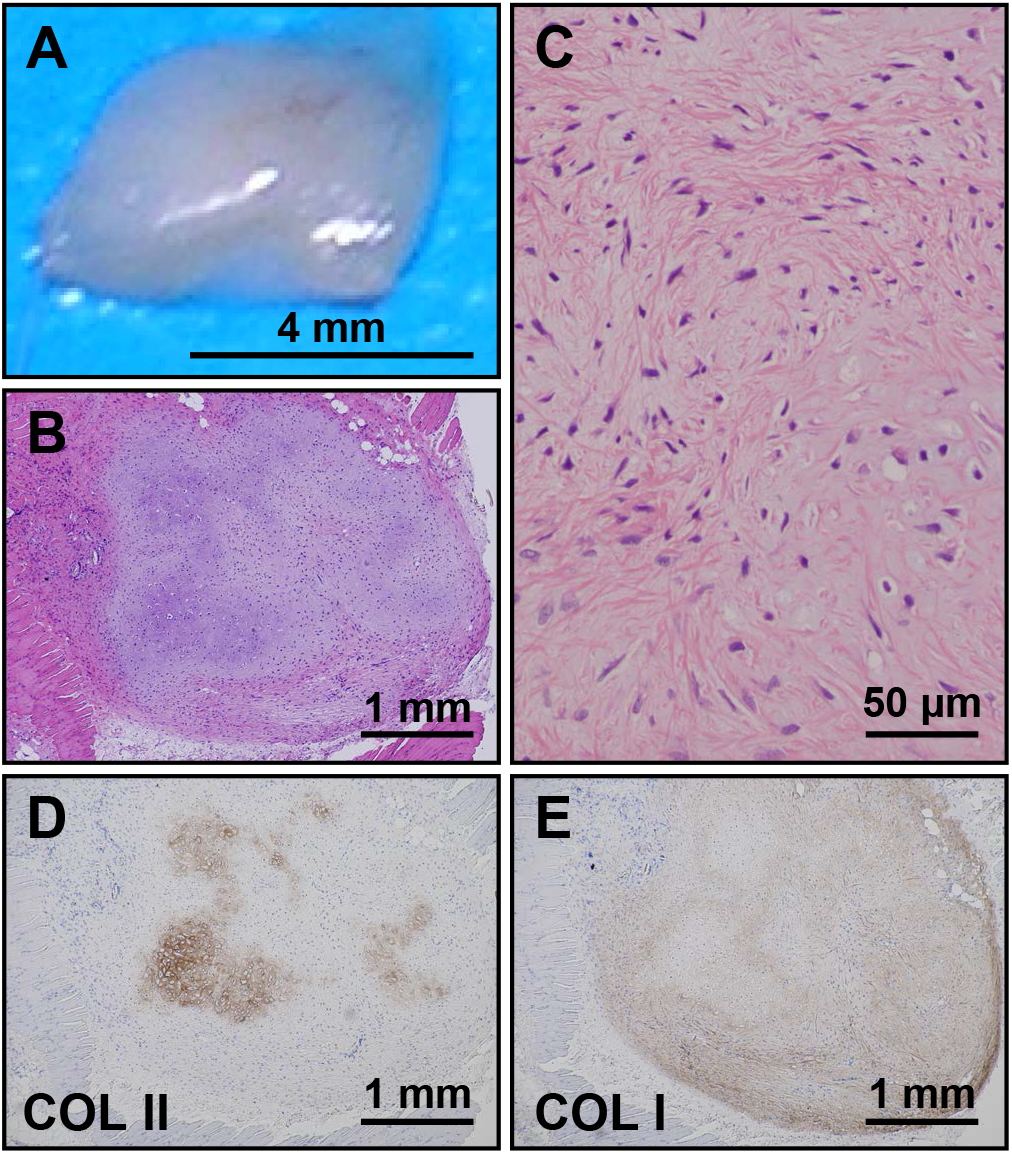
Implantation of P3 cell (PDL 23). P3 (PDL 23) cell-implanted sites were analyzed at 7 weeks. Macroscopic examination showed the flattened and smaller nodules (11 mg in weight) with translucent whitish surface **(A)**. Microscopic examination showed the cartilaginous tissue with tight adhesion to the mouse muscle tissue and reactions of foreign-body type giant cells **(B)**. The nodule was consisted of chondrocytes with spindle nuclei and narrow lacunae and a few hypertrophic cells **(C)** in myxomatous matrix. The matrix was focally positive for COL II **(D)** and diffusely positive for COL I in the cartilage **(E)**.

## Discussion

For the cell-based therapeutic strategies of cartilage and bone formation, the expansion of cultivation are required to gain sufficient number of cells, and to preserve for a long period in recipient, therefore the detailed studies are essential with regards to the characteristics of expanded and/or dedifferentiated chondrocytes in vitro and in vivo.We have previously reported that chondrocytes from epiphyseal cartilage at a primary culture have the ability to regenerate cartilage and to develop bone with marrow cavity through enchondral ossification in vivo [2]. This present study demonstrated the detailed alterations of the human infant epiphyseal chondrocytes during expansions from P0 to P8 cells in monolayer culture and the ability of in vivo chondrogenesis and osteogenesis by implantation, and long-term observation after implantation.

The fate of cartilage generated by chondrocytes depended on population doubling levels of chondrocytes in culture. Primarily, implanted chondrocytes need to retain the abilities to generate cartilage with production of type II collagen-positive cartilaginous matrix, and to produce type I-positive perichondrium. In addition to chondrocytes, participation of undifferentiated mesenchymal cells that differentiate to osteoblasts, adipocytes and endothelia is required for bone development through enchondral ossification [1, 5, 6]. In our previous study [2], we showed that perichondrial spindle cells participate in bone development as undifferentiated mesenchymal cells after implantation of cultivated chondrocytes, like perichondrial cells in a cartilage anlage of human embryonic bone.

Chondrogenic ability of cultivated chondrocytes is associated with surface markers,gene expression, morphology, synthesis of the type of collagen and capacity of redifferentiation, using human articular and other cartilages [7, 8] or animal [9] as cell sources [10-12]. Surface markers include CD26 for the chondrogenic potential of human articular chondrocytes [13] and ratio of CD54 to CD44 as an index of differentiation status [7]. Morphology of chondrocytes changes from a differentiated round to polygonal cell shape to dedifferentiated fibroblast-like phenotype [9, 14]. In gene expression, type-II collagen is essential for chondrocytes capable of generating cartilage [8] and inversely correlated with increasing PDLs [15]. The ratio of type-I and type-II collagen, defined as an index of cell differentiation, is significantly higher at a primary culture and become lower along with expansion [13]. in vivo chondrogenesis is also associated with PDLs of cultivated chondrocytes in rabbit [16, 17], bovine [18], and human fetal and adult articular chondrocytes [19]. Chondrocytes at a primary culture from articular cartilage show mature and well-formed cartilage with extracellular matrix of type-II collagen, however those with chondrocytes with a longer cultivation do not generate cartilage with type-II collagen but form fibrous tissue [16] and almost same results have been presented in chondrocytes from the costal cartilage [17]. Chondrocytes can be not only isolated from cartilage, but also differentiated from somatic stem cells including mesenchymal stem cells and marrow stromal cells [2, 20-23]. Chondrocytes have recently been generated from pluripotent stem cells like other cell types [2, 24-27].

To reverse de-differentiation process during cultivation, several approaches have been developed. Dedifferentiated chondrocytes after a longer cultivated period regain chondrogenic activity by the high-density culture method [28]. To acquire sufficient cell number for cartilage engineering, the co-culture system of in vitro-expanded dedifferentiated chondrocytes with small numbers of primary chondrocyte is useful to re-differentiate the de-differentiated cells and form thicker cartilage tissue with type-II collagen [18]. Wnt/beta-catenin signaling may be involved in maintenance of chondrogenic activity [29, 30]. Cultivation medium is shown to affect cell phenotypes in vitro [31]. In addition to the re-differentiation strategies, the selection approach was employed to isolate functionally active chondrogenic cells with high levels of type-II collagen from dedifferentiated cells or hypertrophic cells with product of type-X collagen for in vivo chondrogenesis [19].

With regard to epiphyseal cartilage, however, the study in concerning cytological descriptions of the isolated culture cells have not been presented and there are a few morphological studies in vivo on the new bone formation by the implantation of chondrocytes isolated from cartilage of rat [32] and mouse [33]. In human epiphyseal chondrocytes, nodular structures resembling mature articular cartilage are formed by primary culture of human fetal epiphyseal chondrocytes for long-term of up to 180 days on hydrogel substrate [34], although the experiment in passage-expansion of cultivated cells was not performed. Our observations in the present studies of epiphyseal chondrocytes showed the morphology in vivo on the regenerated cartilage and new bone formation with marrow cavity by the implantation of expanded cells for short to long period. Large cartilage without ossification needs to be formed by implantation of cultivated cells with higher PDLs by usage of cytokines in future.

Finally, the present studies showed the changes in cytology and gene expression of the infant epiphyseal chondrocytes in a monolayer culture by cell expansion from the P0 to P8 cells. The experiment in vivo by implantation of expanded cells for a short to long periods demonstrated the fate of the epiphyseal chondrocytes and elucidated that the proceeded passage-cells can regenerate the tissue for a short period; however these tissues can’t be preserved for a longer period and absorbed, except those of the P0 and P1 cells. The bone development through enchondral ossification occurred in cartilage by the P0 cells and by the P1 cells with only lower PDLs. The P1 cells with rather higher PDLs form the articular cartilage-like tissue without ossification. These findings would be a help of chondrocyte-based therapy.

## Acknowledgement

We would like to express our sincere thanks to T. Sugiki, H. Takahashi, and Y. Ito for support throughout the work, H. Abe for excellent technical assistances, and E. Suzuki and K. Saito for assistant in preparation of the manuscript.

## Conflict of interest

The authors declare that there is no conflict of interest regarding the publication of this manuscript.

## Reference

1. Rhodin JAG. Cartilage: bone and bone development. Histology: A text and atlas. New York: Oxford University press; 1974. p. 173-220.

2. Nasu M, Takayama S, Umezawa A. Endochondral ossification model system: Designed cell fate of human epiphyseal chondrocytes during long-term implantation. J 7 Cell Physiol. 2015. doi: 10.1002/jcp.24882. PubMed PMID: 25640995.

3. Kabara M, Kawabe J, Matsuki M, Hira Y, Minoshima A, Shimamura K, et al. Immortalized multipotent pericytes derived from the vasa vasorum in the injured vasculature. A cellular tool for studies of vascular remodeling and regeneration. Lab Invest. 2014;94(12):1340-54. doi: 10.1038/labinvest.2014.121. PubMed PMID: 25329003.

4. Dimri GP, Lee X, Basile G, Acosta M, Scott G, Roskelley C, et al. A biomarker that identifies senescent human cells in culture and in aging skin in vivo. Proc Natl Acad Sci U S A. 1995;92(20):9363-7. PubMed PMID: 7568133; PubMed Central PMCID: PMCPMC40985.

5. Gartner L, Hiatt J. Cartilage and bone. Color Textbook of Histology. 3rd ed. Philadelphia: Saunders; 2007. p. 131-56.

6. Rosenberg AE, Roth SI. Bone. In: Mills SE, editor. Histology for Pathologists. 3rd ed. Philadelphia: Lippincott Williams & Wilkins; 2007. p. 76-95.

7. Hamada T, Sakai T, Hiraiwa H, Nakashima M, Ono Y, Mitsuyama H, et al. Surface markers and gene expression to characterize the differentiation of monolayer expanded human articular chondrocytes. Nagoya J Med Sci. 2013;75(1-2):101-11. PubMed PMID: 23544273; PubMed Central PMCID: PMCPMC4345713.

8. Schnabel M, Marlovits S, Eckhoff G, Fichtel I, Gotzen L, Vécsei V, et al. Dedifferentiation-associated changes in morphology and gene expression in primary human articular chondrocytes in cell culture. Osteoarthritis Cartilage. 2002;10(1):62-70. 28 doi: 10.1053/joca.2001.0482. PubMed PMID: 11795984.

9. Kino-Oka M, Maeda Y, Sato Y, Maruyama N, Takezawa Y, Khoshfetrat AB, et al. Morphological evaluation of chondrogenic potency in passaged cell populations. J Biosci Bioeng. 2009;107(5):544-51. doi: 10.1016/j.jbiosc.2008.12.018. PubMed PMID: 419393556.

10. Holtzer H, Abbott J, Lash J, Holtzer S. THE LOSS OF PHENOTYPIC TRAITS BY DIFFERENTIATED CELLS IN VITRO, I. DEDIFFERENTIATION OF CARTILAGE CELLS. Proc Natl Acad Sci U S A. 1960;46(12):1533-42. PubMed PMID: 16590779; PubMed Central PMCID: PMCPMC223078.

11. Mayne R, Vail MS, Mayne PM, Miller EJ. Changes in type of collagen synthesized as clones of chick chondrocytes grow and eventually lose division capacity. Proc Natl Acad Sci U S A. 1976;73(5):1674-8. PubMed PMID: 1064040; PubMed Central PMCID: PMCPMC430362.

12. Grundmann K, Zimmermann B, Barrach HJ, Merker HJ. Behaviour of epiphyseal mouse chondrocyte populations in monolayer culture. Morphological and immunohistochemical studies. Virchows Arch A Pathol Anat Histol. 1980;389(2):167-87. PubMed PMID: 7456325.

13. Diaz-Romero J, Nesic D, Grogan SP, Heini P, Mainil-Varlet P. Immunophenotypic changes of human articular chondrocytes during monolayer culture reflect bona fide dedifferentiation rather than amplification of progenitor cells. J Cell Physiol. 2008;214(1):75-83. doi: 10.1002/jcp.21161. PubMed PMID: 17559082.

14. Marlovits S, Hombauer M, Truppe M, Vècsei V, Schlegel W. Changes in the ratio of type-I and type-II collagen expression during monolayer culture of human chondrocytes. J Bone Joint Surg Br. 2004;86(2):286-95. PubMed PMID: 15046449.

15. Lin Z, Fitzgerald JB, Xu J, Willers C, Wood D, Grodzinsky AJ, et al. Gene expression profiles of human chondrocytes during passaged monolayer cultivation. J Orthop Res. 2008;26(9):1230-7. doi: 10.1002/jor.20523. PubMed PMID: 18404652.

16. Kang SW, Yoo SP, Kim BS. Effect of chondrocyte passage number on histological aspects of tissue-engineered cartilage. Biomed Mater Eng. 2007;17(5):269-76. PubMed PMID: 17851169.

17. Lee J, Lee E, Kim HY, Son Y. Comparison of articular cartilage with costal cartilage in initial cell yield, degree of dedifferentiation during expansion and redifferentiation capacity. Biotechnol Appl Biochem. 2007;48(Pt 3):149-58. doi: 10.1042/BA20060233. PubMed PMID: 17492943.

18. Gan L, Kandel RA. In vitro cartilage tissue formation by Co-culture of primary and passaged chondrocytes. Tissue Eng. 2007;13(4):831-42. doi: 10.1089/ten.2007.13.ft-358. PubMed PMID: 17253927.

19. Wu L, Gonzalez S, Shah S, Kyupelyan L, Petrigliano FA, McAllister DR, et al. Extracellular matrix domain formation as an indicator of chondrocyte dedifferentiation and hypertrophy. Tissue Eng Part C Methods. 2014;20(2):160-8. doi: 10.1089/ten.TEC.2013.0056. PubMed PMID:23758619; PubMed Central PMCID: 13 PMCPMC3910562.

20. Abe-Suzuki S, Kurata M, Abe S, Onishi I, Kirimura S, Nashimoto M, et al. CXCL12+ stromal cells as bone marrow niche for CD34+ hematopoietic cells and their association with disease progression in myelodysplastic syndromes. Lab Invest. 2014;94(11):1212-23. doi: 10.1038/labinvest.2014.110. PubMed PMID: 25199050.

21. Takano T, Li YJ, Kukita A, Yamaza T, Ayukawa Y, Moriyama K, et al. Mesenchymal stem cells markedly suppress inflammatory bone destruction in rats with adjuvant-induced arthritis. Lab Invest. 2014;94(3):286-96. doi: 10.1038/labinvest.2013.152. PubMed PMID: 24395111.

22. Umezawa A, Maruyama T, Segawa K, Shadduck RK, Waheed A, Hata J. Multipotent marrow stromal cell line is able to induce hematopoiesis in vivo. J Cell Physiol. 1992;151(1):197-205. doi: 10.1002/jcp.1041510125. PubMed PMID: 1373147.

23. Sugiki T, Uyama T, Toyoda M, Morioka H, Kume S, Miyado K, et al. Hyaline cartilage formation and enchondral ossification modeled with KUM5 and OP9 chondroblasts. J Cell Biochem. 2007;100(5):1240-54. doi: 10.1002/jcb.21125. PubMed PMID: 17115412.

24. Santostefano KE, Hamazaki T, Biel NM, Jin S, Umezawa A, Terada N. A practical guide to induced pluripotent stem cell research using patient samples. Lab Invest. 2015;95(1):4-13. doi: 10.1038/labinvest.2014.104. PubMed PMID:25089770.

25. De Assuncao TM, Sun Y, Jalan-Sakrikar N, Drinane MC, Huang BQ, Li Y, et al. Development and characterization of human-induced pluripotent stem cell-derived cholangiocytes. Lab Invest. 2015;95(6):684-96. doi: 10.1038/labinvest.2015.51. PubMed PMID: 25867762; PubMed Central PMCID: PMCPMC4447567.

26. De Assuncao TM, Sun Y, Jalan-Sakrikar N, Drinane MC, Huang BQ, Li Y, et al. Development and characterization of human-induced pluripotent stem cell-derived cholangiocytes. Lab Invest. 2015;95(10):1218. doi: 10.1038/labinvest.2015.99. PubMed PMID: 26412498.

27. Higuchi A, Ling QD, Kumar SS, Munusamy MA, Alarfaj AA, Chang Y, et al. Generation of pluripotent stem cells without the use of genetic material. Lab Invest. 2015;95(1):26-42. doi: 10.1038/labinvest.2014.132. PubMed PMID: 25365202.

28. Ahmed N, Gan L, Nagy A, Zheng J, Wang C, Kandel RA. Cartilage tissue formation using redifferentiated passaged chondrocytes in vitro. Tissue Eng Part A. 2009;15(3):665-73. doi: 10.1089/ten.tea.2008.0004. PubMed PMID: 18783319.

29. Usami Y, Gunawardena AT, Iwamoto M, Enomoto-Iwamoto M. Wnt signaling in cartilage development and diseases: lessons from animal studies. Lab Invest. 2016;96(2):186-96. doi: 10.1038/labinvest.2015.142. PubMed PMID: 26641070.

30. Matsushita K, Morello F, Zhang Z, Masuda T, Iwanaga S, Steffensen KR, et al. Nuclear hormone receptor LXRα inhibits adipocyte differentiation of mesenchymal stem cells with Wnt/beta-catenin signaling. Lab Invest. 2016;96(2):230-8. doi: 10.1038/labinvest.2015.141. PubMed PMID: 26595172; PubMed Central PMCID: PMCPMC4731266.

31. Sztiller-Sikorska M, Hartman ML, Talar B, Jakubowska J, Zalesna I, Czyz M. Phenotypic diversity of patient-derived melanoma populations in stem cell medium. Lab Invest. 2015;95(6):672-83. doi: 10.1038/labinvest.2015.48. PubMed PMID: 25867763.

32. Thyberg J, Moskalewski S. Bone formation in cartilage produced by transplanted epiphyseal chondrocytes. Cell Tissue Res. 1979;204(1):77-94. PubMed PMID: 527023.

33. Moskalewski S, Malejczyk J. Bone formation following intrarenal transplantation of isolated murine chondrocytes: chondrocyte-bone cell transdifferentiation? Development. 1989;107(3):473-80. PubMed PMID: 2612374.

34. Reginato AM, Iozzo RV, Jimenez SA. Formation of nodular structures resembling mature articular cartilage in long-term primary cultures of human fetal epiphyseal chondrocytes on a hydrogel substrate. Arthritis Rheum. 1994;37(9):1338-49. PubMed PMID: 7945499.

